# CoGe LoadExp+: A web-based suite that integrates next-gen sequencing data analysis workflows and visualization

**DOI:** 10.1101/118802

**Authors:** Jeffrey W. Grover, Matthew Bomhoff, Sean Davey, Brian D. Gregory, Rebecca A. Mosher, Eric Lyons

## Abstract

To make genomic and epigenomic analyses more widely available to the biological research community, we have created LoadExp+, a suite of bioinformatics workflows integrated with the web-based comparative genomics platform, CoGe. LoadExp+ allows users to perform transcriptomic (RNA-seq), epigenomic (bisulfite-seq), chromatin-binding (ChIP-seq), variant identification (SNPs), and population genetics analyses against any genome in CoGe, including genomes integrated by users themselves. Through LoadExp+’s integration with CoGe’s existing features, all analyses are available for visualization and additional downstream processing, and are available for export to CyVerse’s data management and analysis platforms. LoadExp+ provides easy-to-use functionality to manage genomics and epigenomics data throughout its entire lifecycle and facilitates greater accessibility of genomics analyses to researchers of all skill levels. LoadExp+ can be accessed at https://genomevolution.org.

## Background

As advanced next-generation sequencing (NGS) technologies become more powerful and affordable, a growing number of researchers find themselves in a position to answer genome-scale biological questions using both existing and newly generated data. However, despite rapid technological advances, the knowledge and computational resources needed to analyze and interpret multiple large genomic datasets create bottlenecks for scientific discovery. While initiatives exist to provide high performance computing resources (e.g., CyVerse, XSEDE) [1, 2] to life scientists, these resources require computational expertise to exploit. Researchers in the life sciences without this expertise would benefit from a platform focused on NGS data and genomics analyses that integrates data management, analysis, and visualization tools into a single user-friendly interface. Such a platform also benefits more advanced users by simplifying the process of analyzing, visualizing, managing, and distributing data with collaborators.

We have addressed these needs with the creation of LoadExp+, an addition to the comparative genomics platform, CoGe (https://genomevolution.org) [3]. LoadExp+ is a web portal through which numerous genomic and epigenomic analyses can be conducted. These analyses include RNAseq, whole-genome bisulfite-sequencing (BS-seq), ChIPseq, SNP identification, and population genetics calculations. The collection of these tools in one location, integration with an advanced genome browser, and the use of CoGe’s user-friendly interface present advantages over other web-based bioinformatics platforms for the life sciences. For advanced users, there is also an REST API available for programmatic access to data integrated through LoadExp+ (https://goo.gl/Pf4xjf).

## Results

### Web Interface

LoadExp+ is accessible from CoGe’s main page from the menu bar under “Tools” or the User Data page by clicking “Load Experimental Data.” It is intuitive to navigate, with mouse-over descriptions and conspicuous links to documentation for each tool. Users begin their experimental set-up by selecting data to load from their local computer, specifying a web address (HTTP or FTP), inputting NCBI Short Read Archive (SRA) accession numbers, or selecting files from the user’s CyVerse Data Store (**Figure 1**). Options for LoadExp+’s integrated programs, otherwise accessible as command-line arguments, are then presented as checkboxes or entry fields. When setting up their desired analysis users must provide minimal metadata by naming and describing their experiment, providing version information, and indicating the source of their data.

**Figure 1.**
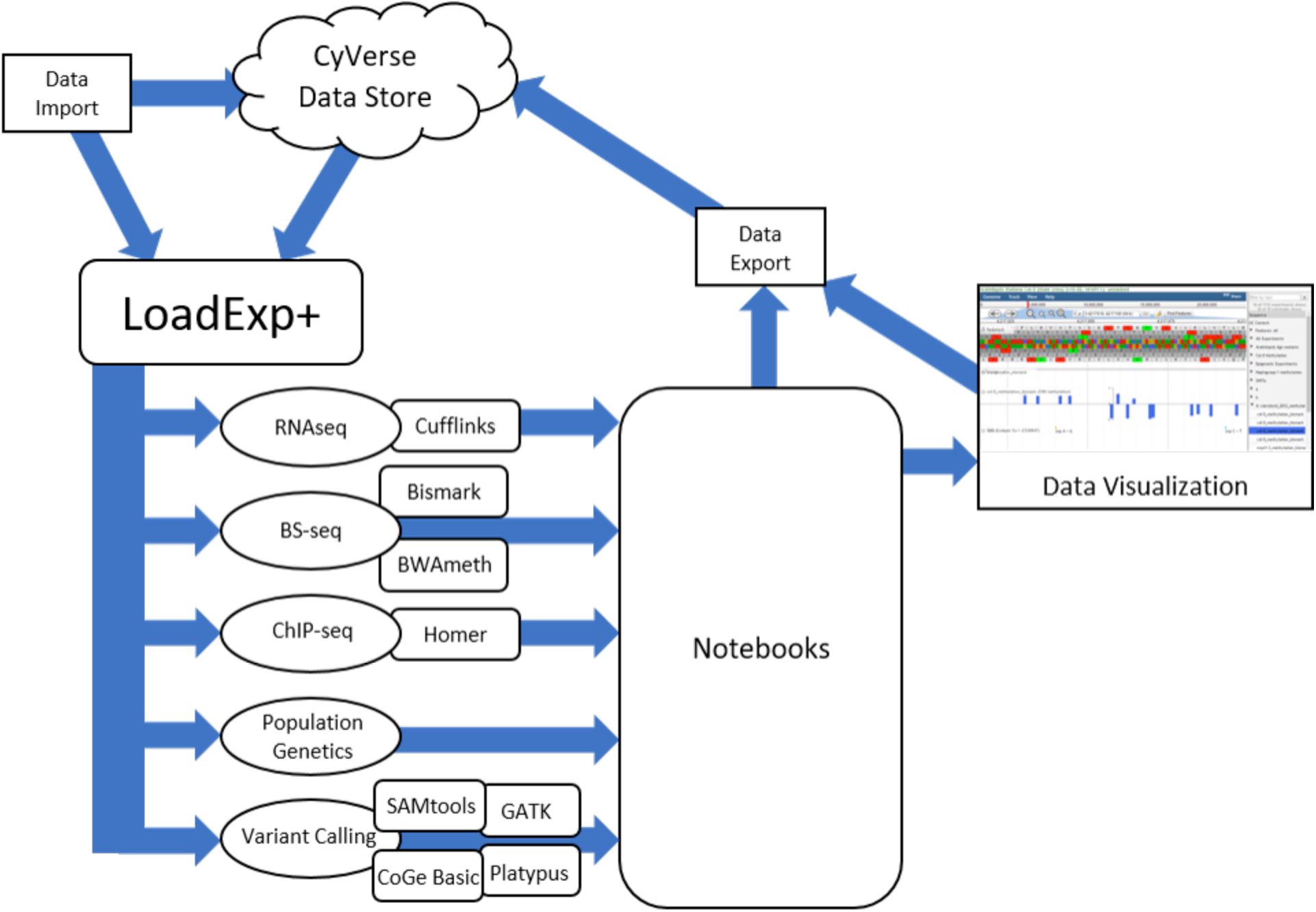
Schematic Representation of LoadExp+ Workflow. Data to be used in any workflow may be imported directly from a user’s local machine, a remote address by HTTP/FTP, from SRA accession number, or from the CyVerse Data Store. After import LoadExp+ allows users to choose which bioinformatics workflows to run. After the desired analyses have finished, experimental data can be organized into “Notebooks” and viewed in CoGe’s integrated genome browser, EPIC-CoGe.

Before submitting their desired analysis users can choose to have the resulting files automatically organized into a new or existing notebook (**Supplemental Figure S1 and S2**), and have an email sent to them when their analysis is complete. After submitting their analysis users are automatically presented with a window in which to monitor its progress as well as a persistent link that they can use to return to monitor the workflow’s progress. Users can also navigate to their “My Data” page to check the progress of submitted LoadExp+ workflows under “Analyses.” Results from the user’s analysis are available for viewing and further analysis in the EPIC-CoGe genome browser (see below). All files, including intermediate files, generated from the analysis can also be downloaded from the Experiment View page (**Supplemental Figure S3 and S4**) or exported to the CyVerse Data Store. These files are also available through a RESTful API following completion of the run.

### Supported Data Formats and Metadata Attribution

LoadExp+ supports the input of data in a variety of formats, including raw sequencing reads (FASTQ, SRA), compressed data (.gz, .bz2, .zip), or preprocessed data in the form of alignments (BAM, SAM), polymorphism data (VCF, GVCF), and quantitative data (CSV/TSV, BED/BEDGraph, GFF/GTF, WIG, BigWig). These data can be uploaded directly from the user’s computer, retrieved from a remote server (FTP/HTTP), imported from their CyVerse Data Store (recommended for files > 1GB), or retrieved from NCBI’s Short Read Archive by accession or project number. Transferring data between CoGe and the CyVerse Data Store uses iRODS [4] for secure, high-performance, parallel file transfers.

Users are prompted to supply metadata which are captured for every experiment, including the name, description, type, and source of the data. Additionally, the analysis program(s), parameters, and the options chosen at run-time are captured automatically and are immutable. After an experiment has been loaded, researchers can add additional metadata including text-based descriptions, web links, and images (**Supplemental Figure S4**).

### Integrated Analysis Workflows

LoadExp+ is a unified platform encompassing a variety of popular and powerful genomic and epigenomic analyses. The following NGS workflows are currently available in LoadExp+: RNA-seq, whole-genome BS-seq, ChIP-seq, SNP identification, and population genetics analysis. More detailed information for each analysis workflow is collected in **Table 1**. A schematic view of LoadExp+ workflows can be seen in **Figure 1** and a full list of all tools (with benchmarking data) is provided in **Supplemental Table S1**. Many workflows share usage of the various aligners integrated into LoadExp+ including GSNAP [12], Bowtie2 [13], TopHat2 [14], or HISAT2 [15]. Sequencing reads can be trimmed by either CutAdapt [16] or Trim Galore! [17].

**Table 1.** Workflow Summary

| LoadExp+ Workflow | Analysis Tool(s) | Detailed Workflow Information (CoGe Wiki) | Visualization (Browser Track) | Analysis |
| --- | --- | --- | --- | --- |
| RNAseq | Cufflinks [5] | https://goo.gl/KCBtbH | Alignment, Read Depth (bar plot), FPKM (bar plot) | FPKM Differential expression |
| Bisulfite-seq | Bismark [6]<br>BWA-meth [7] | https://goo.gl/2eegqK | Alignment, Percent Methylation (bar plot) | Whole-genome DNA methylation |
| ChIPseq | HOMER [8] | https://goo.gl/aFV2T5 | Peaks (bar plot) | Protein-DNA Interactions |
| Variant calling | SAMtools [9], Platypus [10], GATK [11], CoGe Basic | https://goo.gl/GIQGu5 | SNP Genome Browser Track | SNP Discovery |
| Population Genetics | Nucleotide diversity, Watterson's estimator, and Tajima's D | https://goo.gl/j4nNS1 | Custom report | Identification of selection, selective sweeps |

Analyses are dispatched by a job scheduling engine, based on WorkQueue [18]. Users can request a notification email upon completion and monitor the job under the Activity and Data Loading tabs in the My Data page, or with a persistent link displayed on the progress window. All data integrated by users are kept private by default and may be shared with collaborators through the My Data page (**Supplemental Figure S5**). Each LoadExp+ workflow generates output files that can be visualized in the EPIC-CoGe genome browser with a single click. Output files can also be downloaded or exported to CyVerse for users to perform additional analyses.

### Flexible Data Visualization With EPIC-CoGe

We have integrated a customized version of JBrowse [19], called EPIC-CoGe (https://goo.gl/cWX0PB), within CoGe to visualize the data generated by LoadExp+ and other appropriately formatted data. The browser will display any of the thousands of genomes available in CoGe onto which experimental data tracks may be overlaid. Each file generated by LoadExp+ is automatically loaded and made available as selectable tracks for viewing in EPIC-CoGe. Critically, users may simultaneously display their own private data and publicly available data already present in CoGe for the selected genome (**Figure 2**). Users can also mix and match display of different visualization tracks, comparing the results of different NGS workflows as well as data from different labs’ experiments. While browsing a genome of interest and exploring their experimental data users may export results from a genomic region of interest, search quantitative tracks for minima, maxima, or a range of data, and search for SNPs overlapping annotated genomic features. Users can also download complete experimental results, export them directly to their CyVerse Data Store for additional downstream analyses, or access them through CoGe’s RESTful API (https://goo.gl/Pf4xjf).

**Figure 2.**
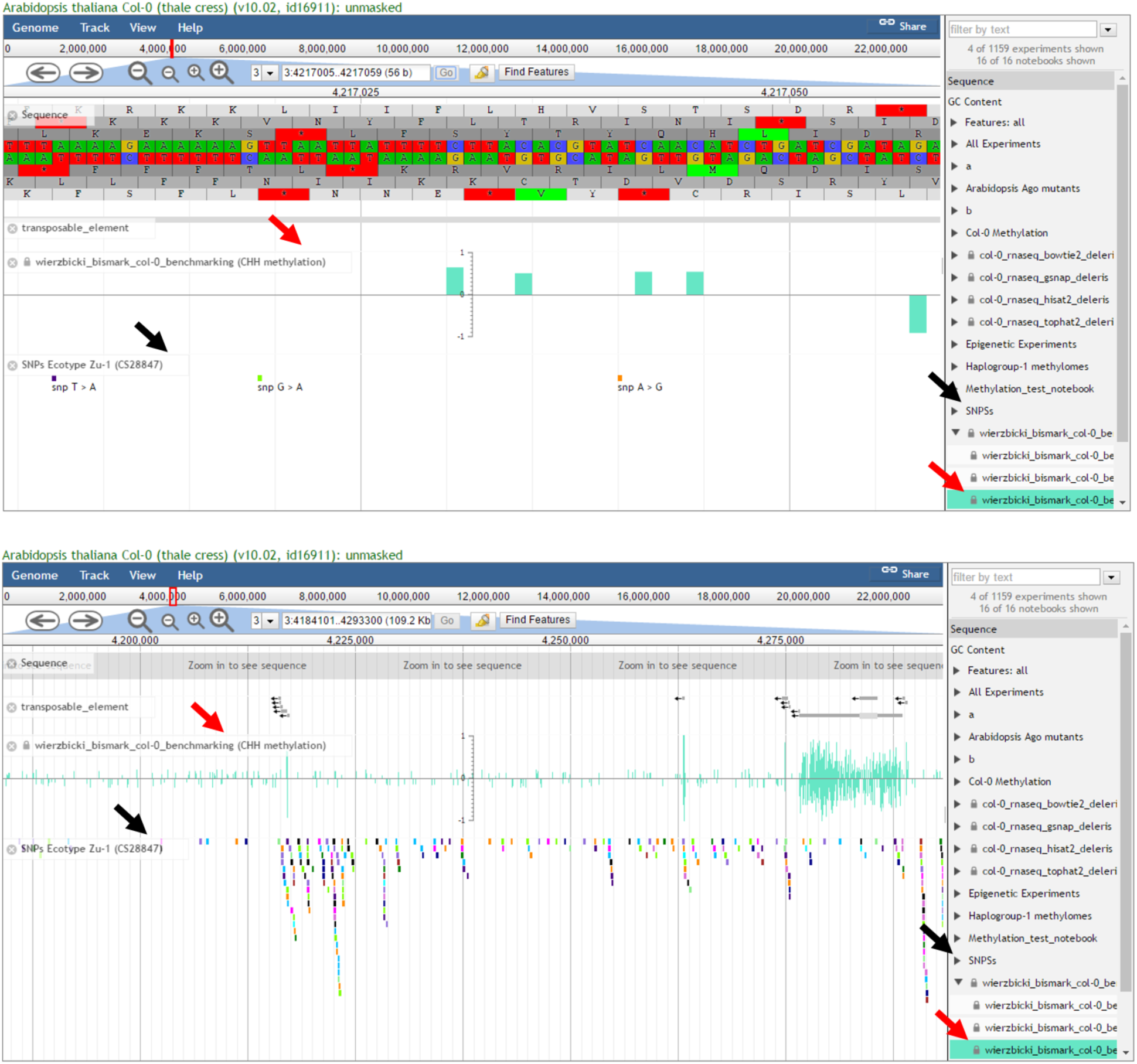
Data Visualization With EPIC-CoGe. At the completion of a LoadExp+ workflow, data are viewed in CoGe’s genome browser, EPIC-CoGe (based on JBrowse). Experiments appear as selectable tracks within each notebook (right side). Shown here are private quantitative data (CHH methylation, red arrows) and public diversity data (SNPs, black arrows) displayed with annotated genomic features (transposable elements) and genomic DNA sequence. The upper and lower images display the same data at different scales to show both specific details (upper) and broad features (lower).

### Benchmarking

To verify that all workflows are functioning as intended we have re-analyzed publicly available data using LoadExp+. In order to simulate the needs of users working in various organisms and with a wide range of data sizes we performed analyses with data from *Arabidopsis thaliana*, a common model organism, *Zea mays*, an agronomically important crop with a large genome, and *Homo sapiens*. We also used both paired ended and single ended reads in our benchmarking. All analyses were completed successfully with no errors, and the options and the run-time are recorded in **Supplemental Table S1-S3**. The slowest-running analysis is often Bisulfite-seq, which is expected because of the computational demands of bisulfite sequencing aligners due to alignment against wild type and C->T converted genomic sequences. The slowest analysis was bisulfite sequencing in *Z. mays* with paired end reads, requiring ã54 hours, while the fastest analysis was RNAseq from *Arabidopsis thaliana*, needing only 36 minutes to complete. These benchmarks represent a general expectation of run times, but users should note that their precise run-time will also vary based on server load.

### CoGe API

Data stored in CoGe, including genomes or the outputs and intermediate files generated by LoadExp+ analyses, are also accessible in a programmatic fashion. By using CoGe’s RESTful [20] application programming interface (API) (https://goo.gl/Pf4xjf) developers can integrate CoGe’s web services into their own websites or analysis pipelines. This extends CoGe and LoadExp+ beyond use by individual researchers by making them available as web services to be utilized by developers.

## Discussion

LoadExp+ allows users to upload, analyze, and visualize a variety of public and private NGS data using CoGe’s web-accessible graphical user interfaces (GUIs) and application programming interfaces (APIs). The streamlined user interface is designed to allow novice users to quickly move from data analysis to visualization, and allows more experienced users to customize their analyses and make them available to collaborators. Data generated, and analyses performed, using LoadExp+ may be kept private, shared with collaborators, or made fully public. Currently, LoadExp+ supports RNA-seq, ChIP-seq, whole-genome bisulfite sequencing (BS-seq), variant analysis (SNP-calling), and population genetics analyses using any genome within CoGe. Users are able to upload genomes for organisms not already present in CoGe, new versions of genomes, and gene model annotations, extending the usefulness of LoadExp+ to any organism with a sequenced genome. CoGe is powered by CyVerse (www.cyverse.org) [1], providing access to high-performance, secure systems for computational scalability and interoperability. LoadExp+ offers a unified approach to NGS data analysis for both novice and experienced users.

LoadExp+ allows users to benefit from CoGe’s CyVerse integration in various ways. The most important of which is federated user identity management, allowing the seamless transfer of large files from a user’s CyVerse Data Store to CoGe in order to perform analyses using any of LoadExp+’s workflows. Data transfers between the Data Store and LoadExp+ are multithreaded and proceed in an automated fashion after submitting a job through the LoadExp+ GUI. Additionally, files that are generated through LoadExp+ analyses can be easily exported to the CyVerse Data Store as well, making them accessible to the bioinformatics applications available through the rest of CyVerse, including the CyVerse Discovery Environment for managing and running additional analyses, and Atmosphere, for applications that require on-demand cloud computing.

LoadExp+ distinguishes itself from similar web-based tools due to its integration with CoGe and easily accessible workflows for numerous genomic and epigenomic analyses. While tools such as Galaxy [21] also combine a user-user friendly interface with high performance computing resources, at the time of publication it is not possible to perform the same analyses available through LoadExp+ on any of the publicly accessible Galaxy servers. Users that wish to perform bisulfite sequencing, in particular, are unserved by public Galaxy installations. In order for users to perform these or other analyses not currently part of a public Galaxy installation they will need to set up their own and install their desired workflows, negating the user-friendly nature of the platform. Many epigenomics analysis tools are also available through the CyVerse Discovery Environment [1]. However, because Discovery Environment applications are integrated by various users, usually for a specific purpose, not all options for the applications may be available. In LoadExp+ we have tried to make visible the most relevant options for each analysis, but users can contact our active development team in the event that an option they desire is not available.

Additionally, LoadExp+ leverages CoGe’s automatic integration of data into our advanced genome browser, EPIC-CoGe. Once data loading or workflows through LoadExp+ are complete, output datasets are automatically available for visualization and additional analyses as tracks in EPIC-CoGe, CoGe’s integrated genome browser which is based on JBrowse [19]. EPIC-CoGe allows users to manipulate visualizations of their data in various ways including changing colors, normalizing or rescaling data, or searching for data that overlaps genomic features of interest. Users can also export raw or processed data, or data from genomic regions of interest either to their local machine or to the CyVerse Data Store.

Finally, using LoadExp+ and CoGe simplifies the management of experimental data. Data generated or loaded through LoadExp+ can be made public or kept private, and can be shared with other researchers easily. Data can also be organized into “Notebooks” that can, similarly, kept private, made public, or shared with collaborators. LoadExp+ and CoGe offer a unified approach to genomics and epigenomics data analysis and visualization for novice and experienced users, as well as facilitating collaboration between scientists.

## Conclusions

To fully exploit NGS technologies in biological research we must increase access to NGS analysis tools. We have tackled this problem through the creation of LoadExp+, an integrated suite of NGS workflows for analysis of genomic and epigenomic data within the CoGe platform. These workflows enable users to easily perform a variety of analyses, share their data with collaborators, and visualize their results on a single, web-based platform with an intuitive GUI. While many web-based platforms exist to assist researchers analyzing NGS data, LoadExp+ provides additional features for managing public and private data, support for many types of NGS data, and seamless integration with the EPIC-CoGe genome browser. We have verified that all LoadExp+ workflows can be run successfully and benchmarked performance with publicly available data (**Supplemental Tables S1-S3**).

Our primary motivation is to lower the barrier of entry into genomics research, but the convenience of LoadExp+ genomics workflows makes their use an attractive option to researchers of all levels. In the time that LoadExp+ has been available, over 250 experiments are loaded each month by researchers (~8 per day), demonstrating the usefulness of this platform.

## Methods

### Benchmarking

Benchmarking was performed using standard options for each workflow and publicly available data (**Supplemental Tables S1-S3**). Analyses were performed one at a time to avoid multiple concurrent analyses affecting run-time, however, run-time was still dependent on day-to-day server load. The results from the benchmarking analyses have also been made public in a CoGe notebook (https://goo.gl/dQoEGJ) and are accessible as experiment files tied to their respective genomes. The experiment files generated through benchmarking are also visible as selectable tracks in the EPIC-CoGe browser.

## Availability of Data and Materials

LoadExp+ is freely available on the web at https://genomevolution.org under the MIT open source license (source code on GitHub https://github.com/LyonsLab/coge). LoadExp+ and EPIC-CoGe user interfaces are written in JavaScript and are compatible with modern web browsers. Server-side components are written in PERL and Python.

## Authors’ contributions

JWG and MB integrated, tested, and benchmarked, the workflows. MB and EL contributed code. SD led development of EPIC-CoGe. EL, RAM, and BDG conceived the project and EL and RAM supervised the development of LoadExp+. JWG, EL, and RAM wrote the manuscript. All authors were involved in editing and approving the final version of this manuscript.

## Acknowledgements

We thank CyVerse (NSF DBI-0735191 and DBI-1265383) for sharing best practices and providing data and computational infrastructure for CoGe (NSF DBI-1265383), and Dr. Brent Pedersen and Dr. Devon Ryan for their assistance in developing the methylation analysis workflows.

